# Biological and Transcriptomic Characterization of Pre-haustorial Resistance to Sunflower Broomrape (*Orobanche cumana* W.)

**DOI:** 10.1101/2021.02.17.431739

**Authors:** Dana Sisou, Yaakov Tadmor, Dina Plakhine, Sariel Hübner, Hanan Eizenberg

## Abstract

Infestations with sunflower broomrape (*Orobanche cumana* Wallr.), an obligatory root parasite, constitute a major limitation to sunflower production in many regions around the world. Breeding for resistance is the most effective approach to reduce sunflower broomrape infestation, yet resistance mechanisms are often overcome by new races of the pathogen. Elucidating the mechanisms controlling the resistance to broomrape at the molecular level is thus the most desirable pathway to obtaining long-lasting resistance and reducing yield loss in sunflower. In this study, we investigated broomrape resistance in a confectionery sunflower hybrid with a robust and long-lasting resistance to sunflower broomrape. Visual screening and histological examination of sunflower roots revealed that penetration of the intrusive broomrape cells into the host root endodermis is blocked at the host cortex, indicating a pre-haustorial mechanism of resistance. A comparative RNA-Seq experiment conducted between roots obtained from the resistant cultivar, a bulk of five broomrape resistant lines and a bulk of five broomrape susceptible lines allowed the identification of genes that were significantly differentially expressed upon broomrape infestation. Among these differentially expressed genes, β-1,3-endoglucanase, β-glucanase and ethylene-responsive transcription factor4 (ERF4) genes were identified. These genes were previously reported to be pathogenesis-related genes in other plant species. This genetics investigation together with the histological examinations led us to conclude that the resistance mechanism involves the identification of the broomrape and the consequent formation of a physical barrier that prevents the penetration of the broomrape into the sunflower roots.

## Introduction

Among the plethora of plant pathogens, parasitic weeds are considered a major threat to crops worldwide. Broomrape species (*Orobanche* and *Phelipanche* spp., Orobanchaceae) are obligatory parasitic plants that are particularly damaging to agricultural crops, especially legumes, tobacco, carrot, tomato, and the crop of interest in this study—sunflower (*Helianthus annuus* L.). Sunflower broomrape (*Orobanche cumana* Wallr.) thus constitutes a major constraint on sunflower production in many regions around the globe, including the Middle East, Southeast Europe, Southwest Asia, Spain, and China (Parker 2013). Since broomrape is a chlorophyll-lacking holoparasite, it obtains all its nutritional requirements from the host plant. The parasitism occurs at the host roots, damaging the host development and resulting in significant yield reduction (Eizenberg et al. 2013). Controlling broomrape is a challenging problem, because only a few herbicides are effective against broomrape and, more importantly, because the parasite’s attachment to the host root tissues allows systemic herbicides to move from the parasite into the host (Eizenberg et al. 2009; 2013). Therefore, breeding for resistant varieties is the most efficient and sustainable means to control broomrape in sunflower. Generally, there are three types of host resistance to broomrape, and these parallel the developmental stage of the parasitism. The first, a pre-attachment resistance mechanism, depends on the ability of the host to prevent the attachment of the parasite, including the prevention of parasite germination and the development and low production or release of germination stimulants (Perez-de-Luque et al. 2008), such as strigolactones, from the host roots into the rhizosphere (Xie at al. 2010; Yoneyama et al. 2013). If pre-attachment resistance fails, broomrape seeds will germinate, and the parasites will grow towards the host roots via chemotropism and attach to the roots (Joel and Bar 2013). The second resistance mechanism – known as post-attachment or pre-haustorial resistance (Perez-de-Luque et al. 2008) – is a mechanism inhibiting penetration into the host root cells and the development of the haustorium, thus preventing vascular conductivity between the parasite and the host (Joel and Losner-Goshen 1994). This resistance involves the production of physical barriers (such as thickening of host root cell walls by lignification and callose deposition (Letousey et al. 2007, Echevarrıa-Zomeno et al. 2006; Perez-de-Luque et al. 2008)), which prevents the parasite from establishing a vascular connecton with the host roots. The third, post-haustorial, type of resistence involves the release of a gum-like substance (Perez-de-Luque et al. 2005; 2006) and the production and delivery of toxic compounds (phenolics) by the host. The transfer of these chemical compounds to the parasite prevent or delay the formation of the tubercles that are necessary for stalk elongation and flowering of the parasite (Perez-de-Luque et al. 2006; Lozano-Baena et al. 2007; Eizenberg et al. 2003). To shed light on the basis of the resistance mechanisms in sunflower, it is first necessary to understand the structure of the plant innate immunity system. The first level of the plant immune system is pathogen-triggered immunity (PTI), which is activated by the recognition of pathogen associated molecular patterns (PAMPs). While, over time, pathogens have developed effectors to inhibit the PAMP-activated PTI response, plants in turn have evolved to perceive and counteract these effectors through a second layer of defense, known as effector-triggered immunity (ETI), formerly known as gene-for-gene resistance (Boller and He 2009). The rapid changes in the race composition of sunflower broomrape have led to an ongoing gene-for-gene ‘arms-race’ between breeders and the parasitic weed. the development of *O. cumana*-resistant cultivars usually includes introgression of resistance genes, which are, in many cases, being overcome by the parasite. This resistance breakdown occurs due to the massive use of vertical (monogenic) resistance (Molinero-Ruiz et al. 2015) and can be addressed by the introduction of horizontal (quantitative) resistance genes with the aim to develop a more durable resistance (Perez-Vich et al. 2004; Roman et al. 2002). In this study, we examined the resistance of the confectionery hybrid cultivar ‘EMEK3’ (developed by Sha’ar Ha’amakim Seeds Ltd.), which has high, long-term resistance to sunflower broomrape, with the aim to elucidate – biologically and transcriptomically – the broomrape resistance mechanism, an essential step towards the development of effective sunflower breeding programs.

## Results

### Effect of grafting on the source of the resistance

To determine whether the shoot plays a role in the resistance mechanism, grafting experiments were conducted: a resistant sunflower cultivar (‘EMEK 3’) was cross grafted with a susceptible cultivar (‘DY.3’), and the grafted plants were planted in soil infested with *O. cumana* seeds (20 mg seeds/liter of soil). Self-grafted and non-grafted plants served as controls. All the plants roots of the susceptible variety were parasitized by 402-440 *O. cumana* tubercles and stalks of different sizes, while no parasitism was observed in roots of plants of the resistant cultivars (Fig. 1a,b).

**Fig. 1.**
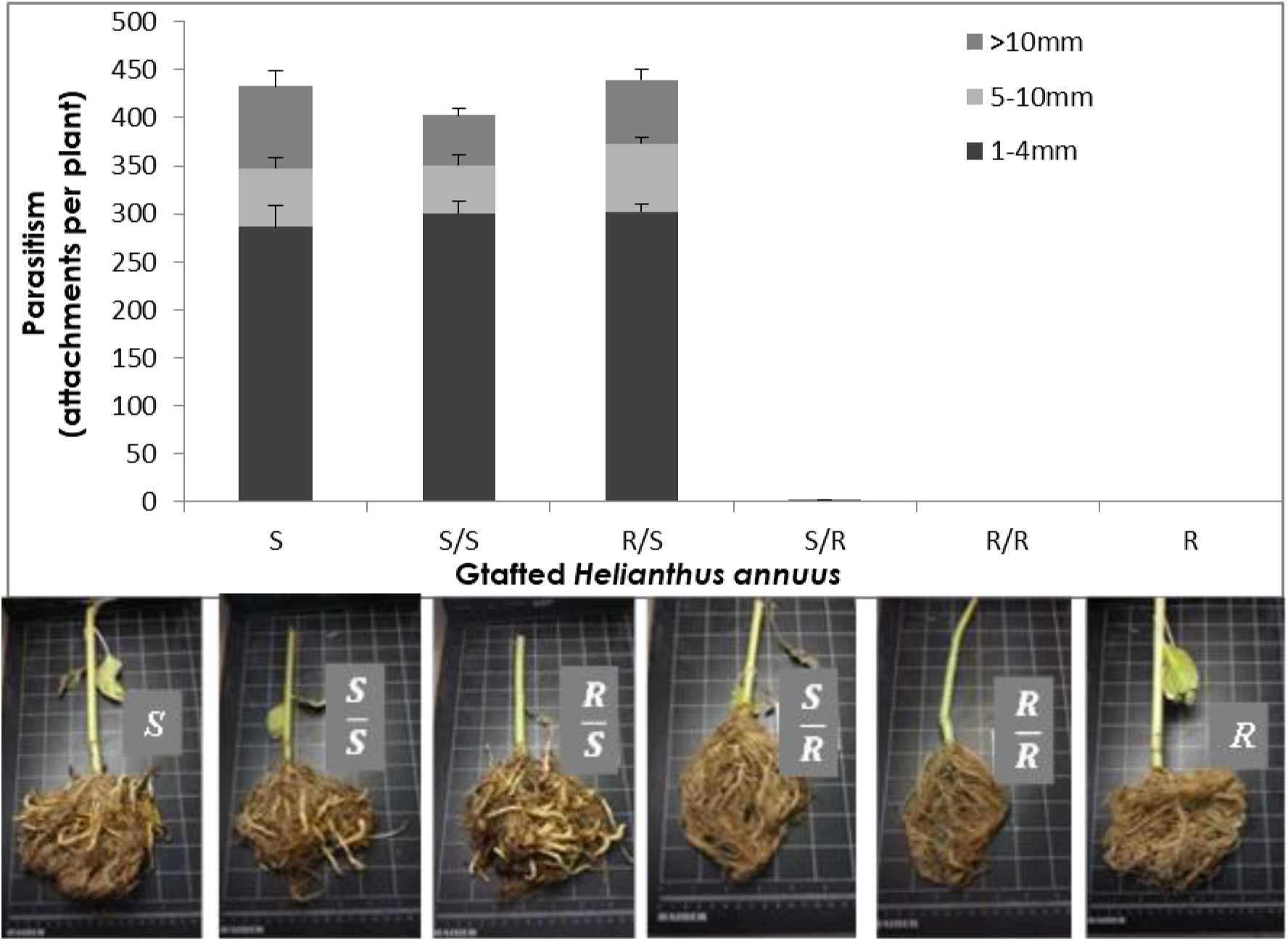
(**a**) Mean number (◻SE) of O. cumana tubercles parasitizing grafted resistant and susceptible sunflower plants. S - non-grafted susceptible sunflower; S/S - self-grafted susceptible sunflower; R/S - resistant sunflower shoot grafted onto susceptible sunflower rootstock; S/R - susceptible sunflower shoot grafted onto resistant sunflower rootstock; R/R - self-grafted resistant sunflower; R - non-grafted resistant sunflower. (b) Grafted and non-grafted sunflower roots 52 days post infestation with O. cumana.

### Sunflower–*O. cumana* incompatibility

Two key parasitism stages were monitored periodically: germination (Fig. 2a) and attachment (Fig. 2b). The germination rate of *O. cumana* seeds was higher (50%) in the presence of roots of the resistant cultivar than in the presence of roots of the susceptible cultivar (39%) (Fig 2a). The first *O. cumana* attachment was observed 11 days after infestation in the susceptible cultivar, while no attachments were observed in the resistant cultivar (Fig. 2b). Observation of *O. cumana* seedlings growing together with sunflower plantlets in clear polyethylene bags (PEB) revealed that the development of the *O. cumana* seedlings was arrested after attaching and attempting to invade the roots of the resistant cultivar, thereby preventing parasite establishment. The disruption of the penetration of the parasite into the host roots and the subsequent deterioration of the parasite seedlings was accompanied by a darkening of host and parasite tissues at the penetration point. In contrast, establishment and development of healthy tubercles was observed in the roots of the susceptible cultivar (Fig. 3). The time at which the resistance response is induced most strongly after the attachment of the parasite seedling to the host roots was determined by observing the host-parasite system growing in the PEB system for. The highest number of necrotic *O. cumana* seedlings in the presence of the resistant cultivar roots was observed five days after infestation. At this time point, 44% of the germinated *O. cumana* seedlings that had attached to the resistant variety roots were necrotic, while on the susceptible variety roots only 9% appeared necrotic (Fig. S1). Histological examination showed that the intruding *O. cumana* cells were blocked at the cortex of the resistant variety roots and could not reach the endodermis (Fig. 4c,d). The root endodermal cells and the seedling intruding cells were stained with safranin, indicating the presence of phenolic compounds which prevented the connection of the parasite to the host vascular system and hence the development of the parasite (Fig. 4).

**Fig. 2.**
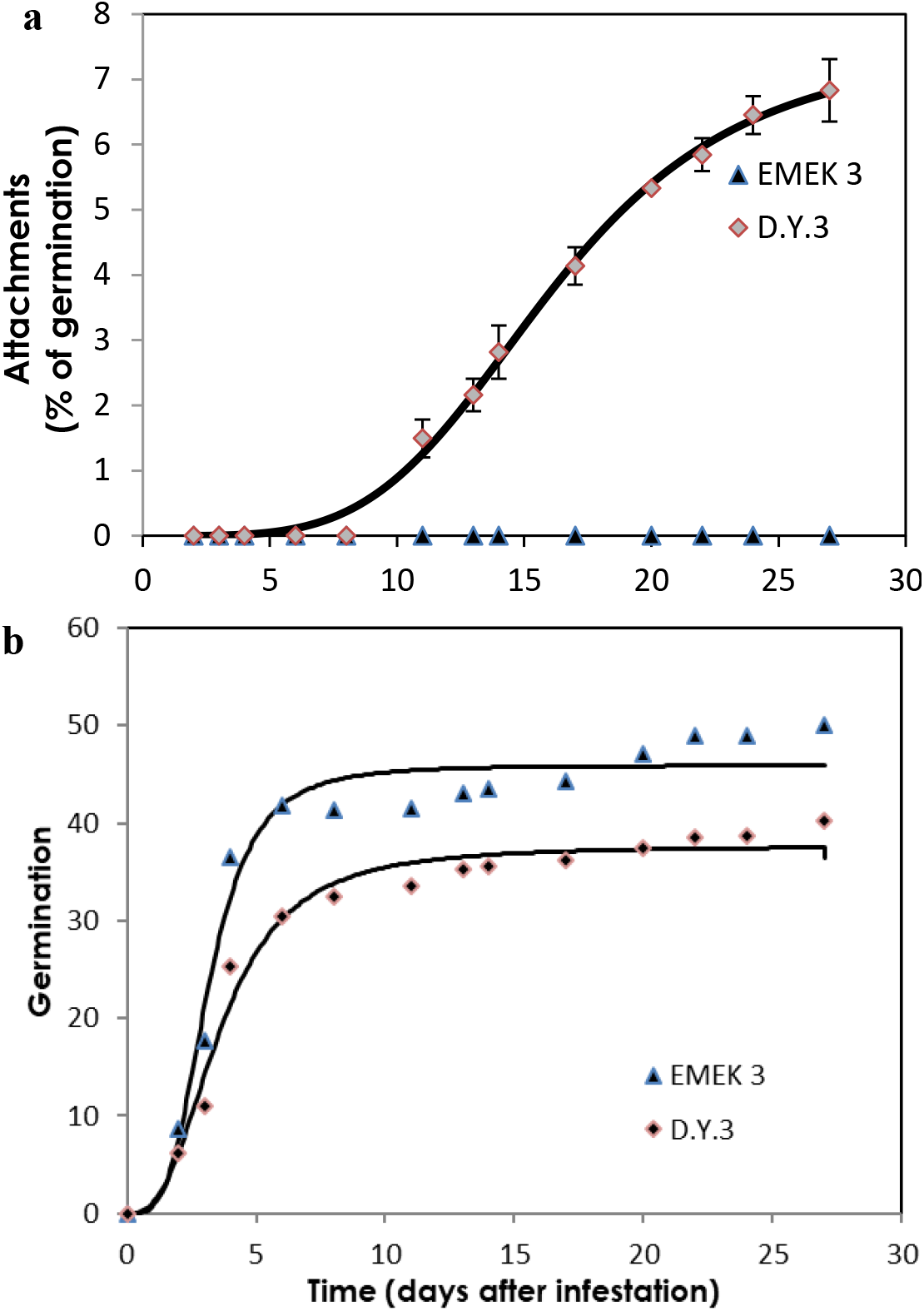
Parasitism dynamics of O. cumana on non-grafted resistant and susceptible sunflower grown in a polyethylene bag system. Attachment (a) and germination (b) of O. cumana in the presence of resistant (‘EMEK3’) and susceptible (‘D.Y3’) sunflower cultivars.

**Fig. 3.**
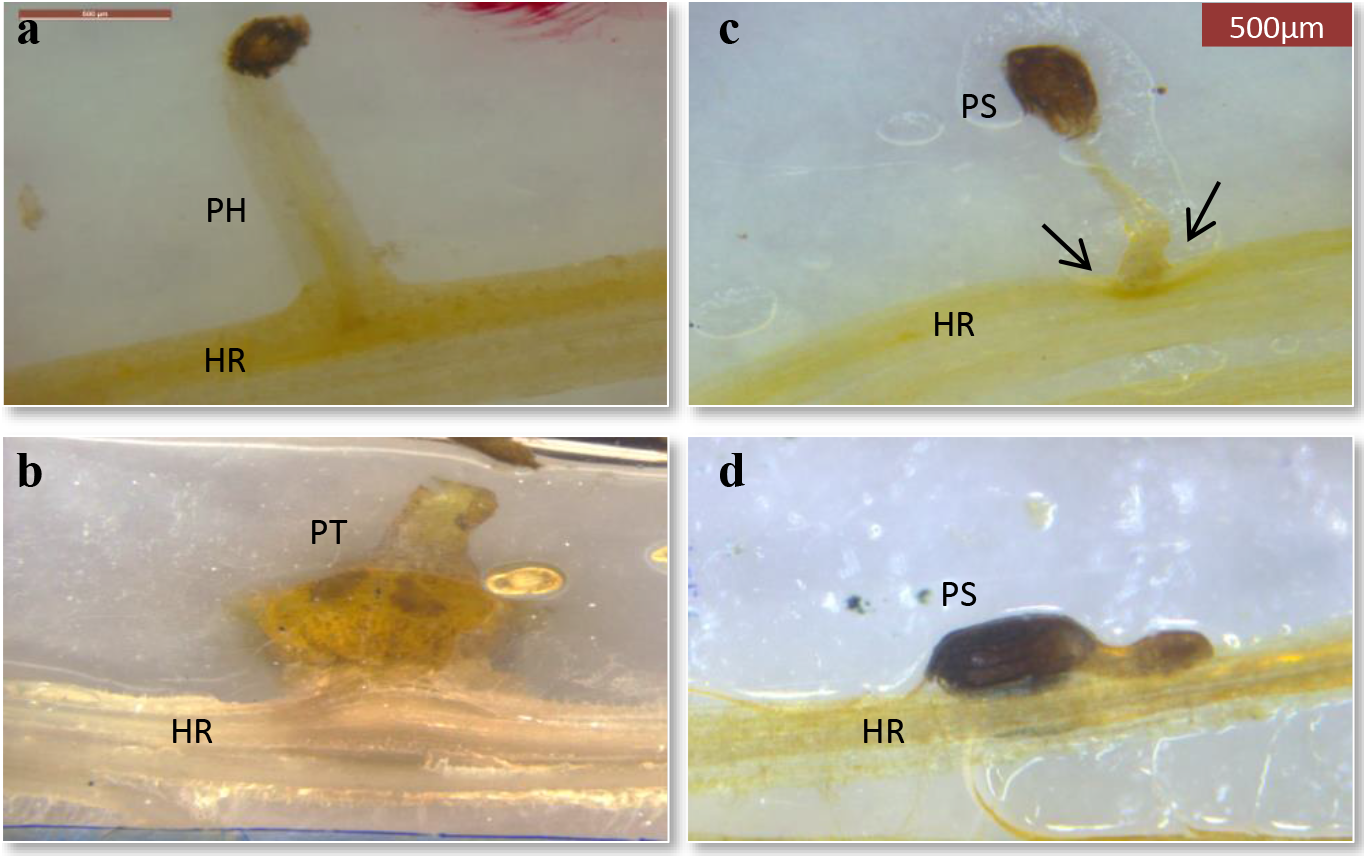
Resistant (c,d) and susceptible (a,b) sunflower roots infested with O. cumana, 10 (a,c) and 21 (b,d) days post inoculation. PH - parasite haustorium; HR – host root; PS – parasite seedling; PT – parasite tubercle.

**Fig. 4.**
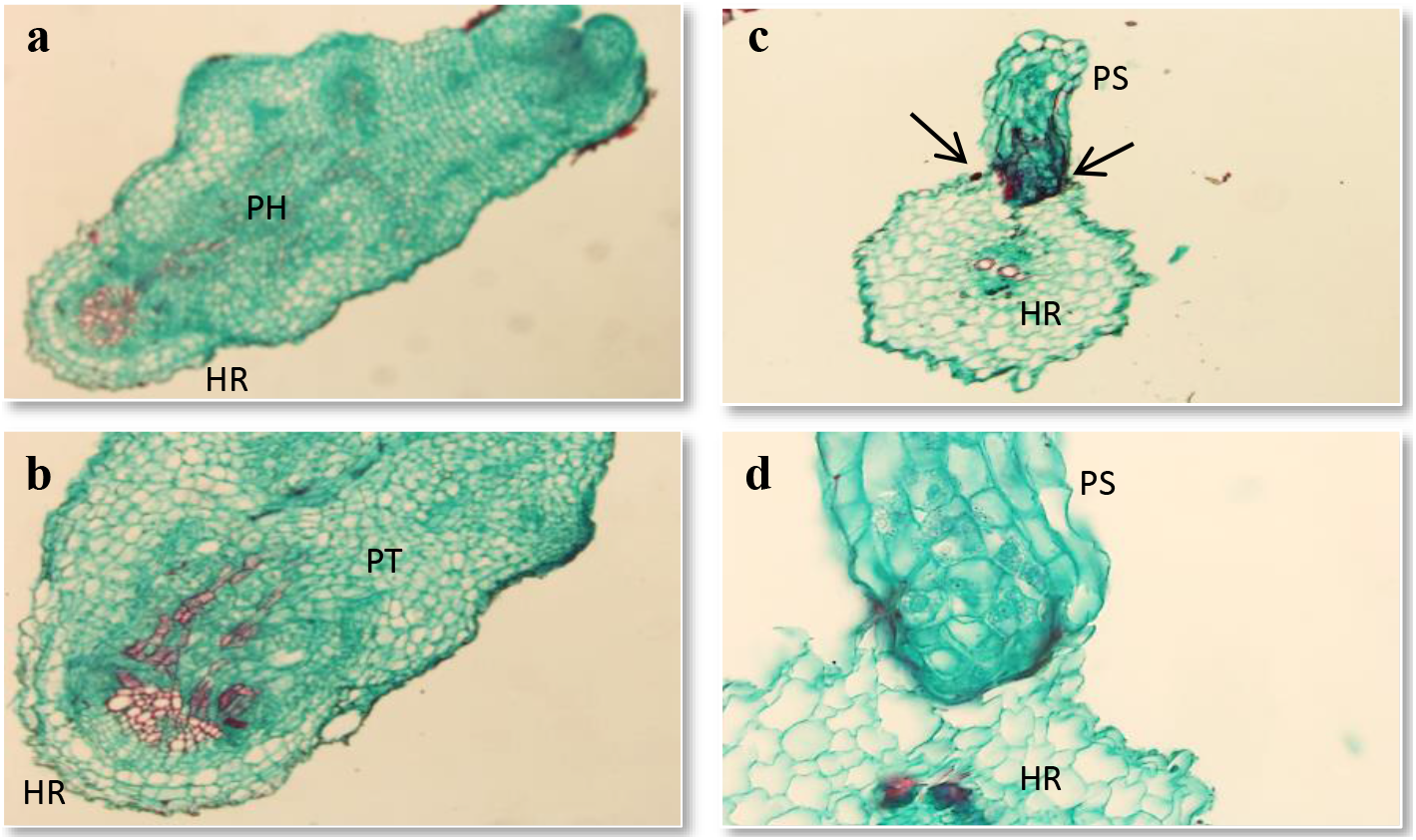
Cross-sections of compatible and incompatible interactions of O. cumana with susceptible (a,b) and resistant (c,d) sunflower roots. PH - parasite haustorium; HR – host root; PS – parasite seedling; PT – parasite tubercle.

### Identification of candidate resistance genes by using RNA-sequencing

Twelve *O. cumana* resistant and susceptible sunflower accessions, which had been used as a genetic source for ‘EMEK3’ breeding, were quantified for *O. cumana* resistance under conditions of artificial infestation in pots held in a greenhouse (25-30 °C). Five accessions showed complete resistance with no attachments on the roots, and seven lines exhibited susceptibility at all *O. cumana* parasitism stages (Fig. S2). Therefore, five resistant accessions and five susceptible accessions were selected to construct a resistant (R) bulk and a susceptible (S) bulk for RNA sequencing (RNA-Seq). Comparative RNA-Seq of *O. cumana* infested and non-infested sunflower roots of the resistant (E) cultivar, the R bulk and the S bulk was used to identify differentially expressed genes (DEGs) associated with sunflower resistance. Twenty-seven barcoded cDNA libraries were constructed and sequenced on two lanes of an Illumina HiSeq 2500 run. A total of 7.4-9.8×10^6^ reads from each library were produced, with an average of 8,648,866.3 reads (Table S1). As our preliminary results showed that the resistance response was strongly induced 5 days after infestation with preconditioned *O. cumana* seeds (Fig. S1), we collected the samples at that time.

Gene Ontology (GO) enrichment analysis performed on 1439 significant DEGs in the resistant cultivar showed that 224 overexpressed genes were significantly enriched [false discovery rate (FDR) < 0.05]. The most enriched term in the *Biological Process* class was “metabolic process” (74%). For the *Molecular Function* and *Cellular Componen*t classes, these terms were “catalytic activity” (67%) and “cell periphery” (14%), respectively (Fig. 5).

**Fig. 5.**
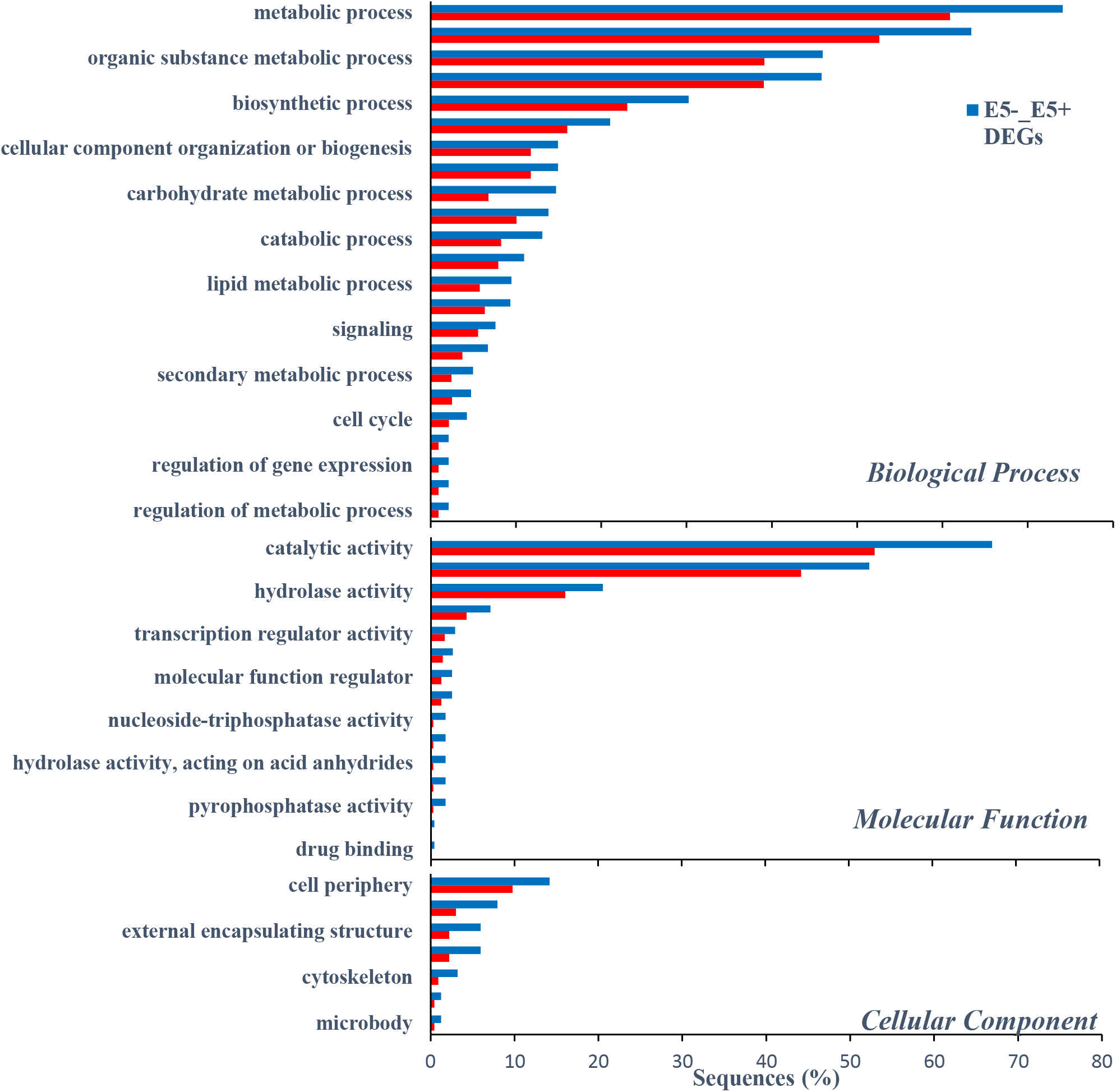
Distribution of enriched GO term for differentially over-expressed genes (Fisher’s Exact Test) for DEGs of the resistant variety EMEK3 (E5-_E5+) compared with GO terms of whole reference predicted gene annotation (HanXRQ). Y-axis represents significant enrichment of GO terms and X-axis shows the frequency of the term.

Out of 1123 and 348 genes that were differentially expressed pre-infestation and 5 days post infestation in the R bulk and ‘EMEK3’ respectively, 37 genes were found to be communal and not differentially expressed in the S bulk (Fig. 6a). To exclude genes that are not related to broomrape infestation, we cross-compared the DEGs of non-infested samples collected on the infestation day and at 5 days post infestation: 47 genes were found to be communal to R bulk and ‘EMEK3’ (Fig. 6b). These 47 genes were then cross-compared with the 37 genes previously mentioned. Two genes were found to be communal and therefore discarded (Fig. 6c). Hence, 35 genes were classified as related to the resistance response (Fig. 6c). Thereafter, we cross-compared the genes that were differentially expressed in ‘EMEK3’ between all treatments: since there was a five-day difference between sampling dates, we assumed that some of the DEGs were not related to the resistance response (i.e., regulatory genes). Therefore, we focused on the 44 genes that were differentially expressed pre-infestation and at the time corresponding to 5 days post infestation with and without *O. cumana* (Fig. 6d). Finally, these 44 DEGs were cross-compared with the 35 DEGs communal to R bulk and ‘EMEK3’ during the resistance response; three genes were found to be mutual (Fig. 6e). These three genes were annotated to the sunflower genome and were identified as β-glucanase, β-1,3-endoglucanase and ethylene-responsive transcription factor 4 (ERF4). The expression levels of the genes encoding β-1,3-endoglucanase and β-glucanase were, respectively, 2.49 and 2.5 times higher in ‘EMEK3’ roots. The expression level of the gene encoding ERF4 was 2.97 times lower in ‘EMEK3’ roots 5 days after the infestation with *O. cumana* (Fig. 7).

**Fig. 6.**
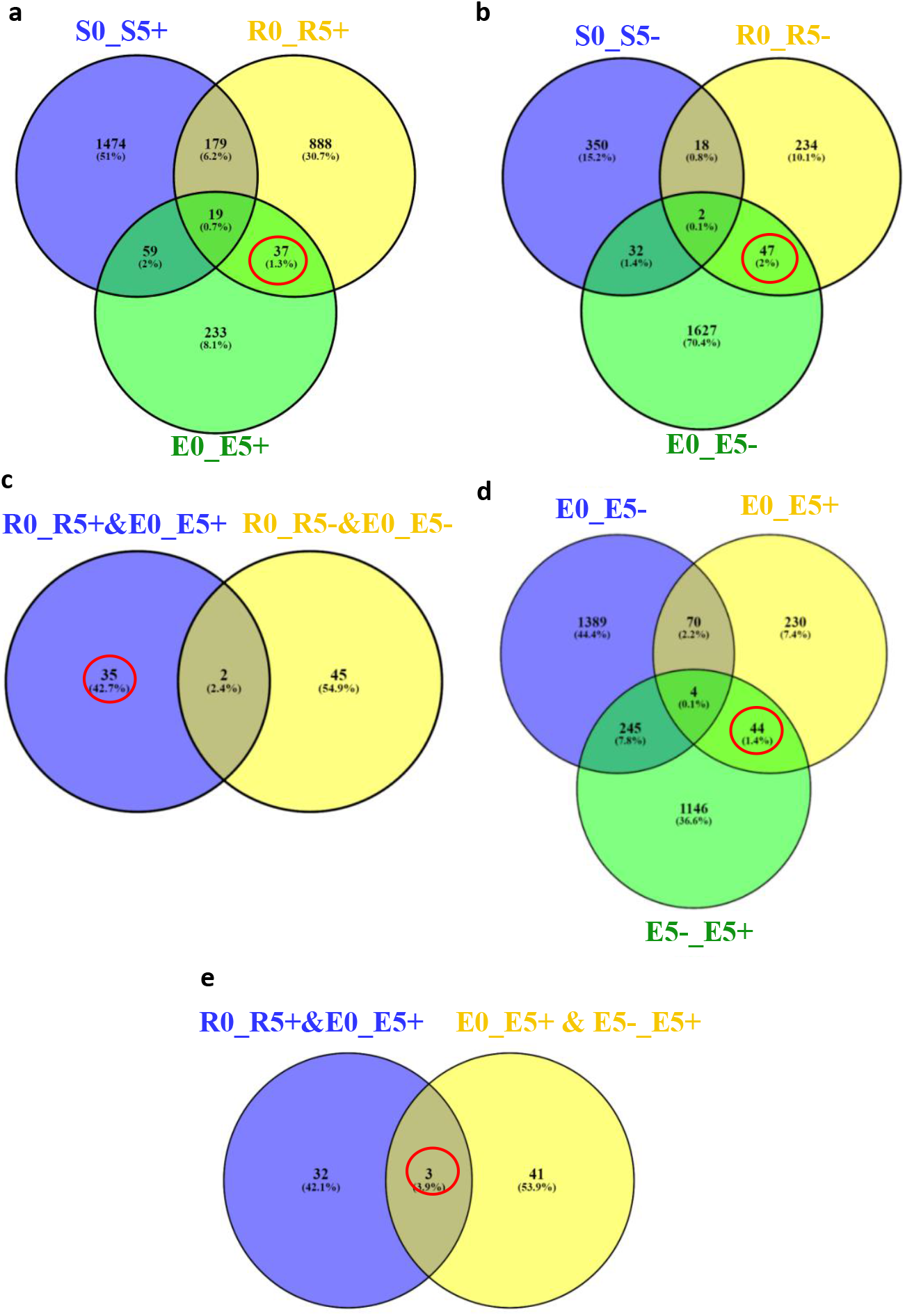
Venny diagram of the DEG between the R bulk (R), S bulk (S) and EMEK3 (E), pre inoculation (0), five days post inoculation with *O.cumana* (5+) (**a**) and five days post inoculation without *O.cumana* (5−) (**b**). (**c**) venny diagram of the communal DEG of **a** and **b**. (**d**) the DEG in EMEK3 pre and 5 days post inoculation with or without *O.cumana*. (**e**) venny diagram of the communal DEG of c and d

**Fig. 7.**
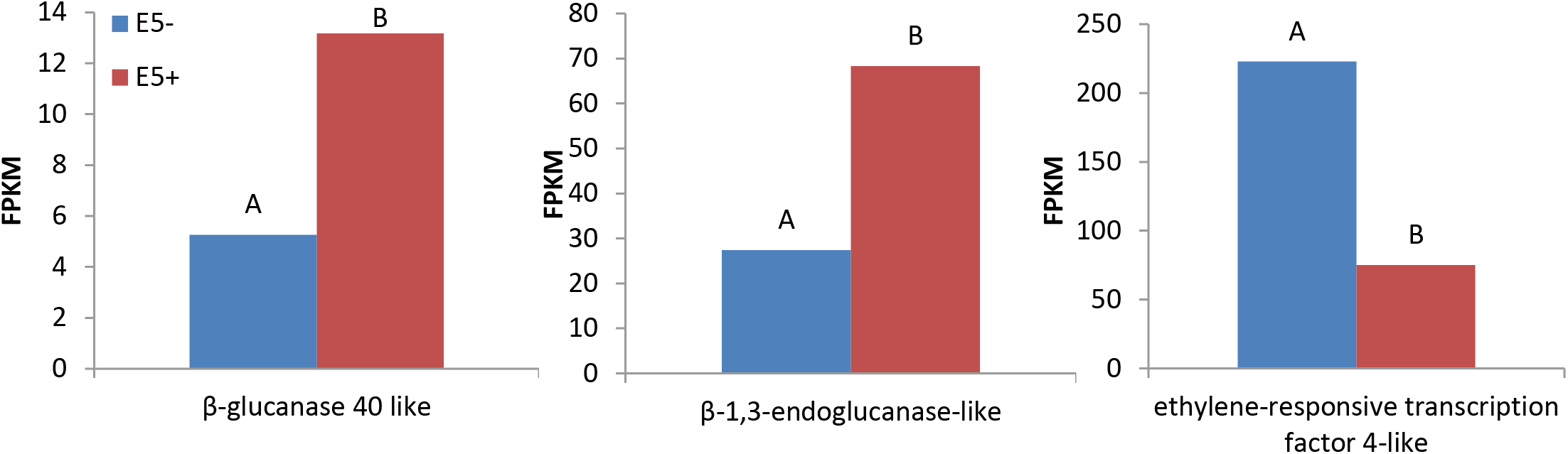
expression levels of β-glucanases, β-1,3-endoglucanase and ethylene-responsive transcription factor 4-like in EMEK3 roots 5 days post infestation of infested (E5+) and non infested (E5+) roots

## Discussion

A variety of strategies comprising the host defense response at the early stages of the parasite life cycle have been described in a number of studies, namely, lignification and subarization of host cell walls (Echevarria-Zomeno el al. 2006), accumulation of callose, peroxidases, and H2O2 in the cortex and protein cross-linking in the cell walls (Perez-de-Luque et al. 2005; 2006), phenylalanine ammonia lyase (PAL) activity and high concentrations of phenolic compounds in the host roots (Goldwasser et al. 1999; 2000; Yang et al. 2017) and degeneration of tubercles after establishment (Eizenberg et al. 2003). Determining the phenological stage at which the incompatibility occurs is crucial for understanding the resistance mechanism and the molecular basis that governs it. To this end, we set up an observation system, based on transparent PEBs, that enabled us to follow the sunflower–*O. cumana* interaction on an ongoing basis and thereby to overcome the difficulty of detecting the exact time at which response is maximal. Our observations revealed that the resistance response began in the early stages of the parasite life cycle, while the parasite was attempting to attach and penetrate into the host roots (Fig. 2–4). Blocking of the penetration attempt was accompanied by necrosis of parasite and host tissues in the penetration area, suggesting a pre-haustorial mechanism of resistance (Perez-de-Luque et al. 2005). Our PEB system showed a markedly high rate of broomrape seedling death in the presence of resistant cultivar roots at 5 days post infestation (Fig. S1). This high death rate was attributed to the prevention of the penetration into the host roots and hence the establishment of the vascular connections that are vital for the parasite seedlings. We thus confirmed by histological methodologies that the parasite intrusion is blocked in the host cortex before the parasite can reach the host endodermis. The endodermal cells in the vicinity of the intrusive broomrape cells in the penetration area were colored with safranin, indicating the involvement of lignin and other phenolic compounds in the host response (Fig. 4). Indeed, suberization, lignification and cell wall thickening have previously been ascribed to the sunflower defense response to *O. cumana* (Gordon Ish-Shalom 1989; Panchenko and Antonova 1974; Echevarria-Zomeno 2006; Jorrin 1996; Yang (2006). We excluded the possibility that the host shoot is involved in the resistance response by grafting susceptible sunflower scions onto resistant rootstocks and vice versa. The resistant cultivar rootstocks conferred resistance on the susceptible scions, but the susceptible rootstocks were parasitized with *O. cumana* regardless of whether the grafted scions were resistant or susceptible (Fig. 1). Similar results have been obtained for the resistance of tomato (*Solanum lycopersicum*) to several broomrape species, namely, *P. aegyptiaca*, *P. ramosa*, *O. cernua* and *O. crenata* (Dor et al. 2010), implying that the resistance response is expressed exclusively in the roots. To elucidate the molecular mechanism that governs the resistance response, we conducted a comparative transcriptome analysis of infested and non-infested resistant and susceptible sunflower roots. The analysis detected 1439 significant DEGs in the roots of the resistant cultivar post infestation. GO and GO enrichment analysis of these DEGs were performed to infer the biological processes and the function of the genes associated with the resistance response, with the ontology analysis revealing a number of overexpressed GO terms (Fig. 5). Importantly, terms associated with the cell periphery (14%), the extracellular region (7.9%), the external encapsulating structure (5.9%) and the cell wall (5.9%) were significantly enriched in the *Cellular Component* category, indicating high activity in these regions. The *Biological Process* category included response to stimulus (21%), cellular component organization (15%) and response to stress (13.8%) (Fig. 5). Finally, a series of Venn diagrams (Oliveros 2015) facilitating cross-comparisons of DEGs in the R bulk, the S bulk and the resistant cultivar ‘EMEK3’ before and after infestation with *O. cumana* identified three genes that were differentially expressed between infested and non-infested sunflower roots of both ‘EMEK3’ and the R bulk (Fig. 6). As a consequence of the infestation, two of these genes, β-1,3-endoglucanase and β-glucanase, were upregulated, and the third gene, ERF4, was downregulated. These findings indicate activation of the plant’s innate immune system, in which the recognition of PAMPs activates a hypersensitive response and accumulation of pathogenesis-related (PR) proteins (Durrant and Dong 2004), such as β-glucanases, which are PR proteins, belonging to the PR-2 family. This family of proteins is believed to play an important role in plant defense responses to pathogen infection (Saboki et al. 2011; Van Loon 1997; Van Loon and Van Strien 1999). Indeed, it has been shown that β-glucanases, which are able to degrade cell wall β-glucan, are involved in resistance to *O. crenata* in pea (*Pisum sativum*) (Perez de luque et al. 2006; Castillejo et al. 2004) and in sunflower resistance to *O. cumana* (Yang et al. 2017). Downregulation of the ERF4 gene post infection should be viewed in the context of the role of the endogenous hormone, ethylene, in regulating defense responses in plants, including the regulation of gene expression during adaptive responses to abiotic and biotic stresses (Anderson 2004). The ERF transcription factors, which are unique to plants, have a binding domain that can bind to the GCC-box, an element found in the promoters of many defense, stress-responsive, and PR genes (Singh et al. 2002; Kazan 2006). Just as there is a range of stresses, there are a large a number of ERFs, with many of the ERFs being transcription activators. Indeed, AtERF1, AtERF2 and AtERF5 act as transcriptional activators, although AtERF3 and AtERF4 act as transcriptional repressors for GCC box-dependent transcription in Arabidopsis leaves (Fujimoto et al. 2000). In that context, McGrath et al. (2005) demonstrated that the Arabidopsis *erf4-1* mutant was resistant to *Fusarium oxysporum*, while transgenic lines overexpressing AtERF4 were susceptible, and therefore concluded that AtERF4 negatively regulates resistance to *F. oxysporum.* The downregulation of the ERF4 gene in the roots of the resistant sunflower post *O. cumana* infestation suggests that, as in Arabidopsis, the sunflower response to biotic stress is negatively regulated by ERF4. Furthermore, the recent study of Liu et al. (2020), using bulked segregant RNA-Seq (BSR-Seq), identified ERF as a candidate gene for *O. cumana* resistance in sunflower.

Taken together, the results obtained for the biological characterization combined with those for the genetic characterization provide a comprehensive view of the relations between the resistant cultivar ‘EMEK3’ and *O. cumana*. This broad view allowed us to propose the following resistance mechanism model: After ‘EMEK3’ induces *O. cumana* seed germination, the seedlings’ attachment to the sunflower roots is perceived by PAMPs. Theses molecules set off a PTI response that downregulates ERF and abrogates the suppression of PR genes (including β-glucanase). β-glucanase then breaks down the parasite cell walls, which, in turn, release effectors that trigger the second level of the plant immune response, namely, effector triggered immunity (ETI). As a result, a physical barrier is created by the accumulation of lignin and other phenolic compounds in the penetration area, and the *O. cumana* seedlings fail to establish a connection with the host vascular system, leading to parasite necrosis.

## Materials and Methods

### Plant material and growth conditions

Twelve sunflower breeding accessions, six resistant and six susceptible to *O. cumana*, and the sunflower varieties ‘EMEK3’ (resistant) and ‘D.Y3’ (susceptible) were kindly provided by Sha’ar Ha’amakim Seeds Ltd (Sha’ar Ha’amakim, Israel). *O. cumana* inflorescences were collected from an infested sunflower field in northern Israel in 2012. The seeds were separated from the capsules using 300-mesh sieves and stored in the dark at 4 °C prior to use. The germination rate of these *O. cumana* seeds at 25 °C was 85%.

### Preconditioning of *O. cumana* seeds

Preconditioning was performed under sterile conditions. The seeds were surface sterilized for 2.5 min in ethanol (70%), followed by10 min in sodium hypochlorite (1%), and then rinsed 5 times with sterile distilled water and dried for 2 h in a laminar airflow cabinet. The dried seeds were spread on a 5.5-cm diameter glass-fiber filter paper discs (Whatman #3, Whatman International Ltd., Maidstone, England) that had been wetted with 600 µl of sterile distilled water. The discs were placed in a sterile 5.5-cm diameter petri dishes. The petri dishes were sealed with Parafilm and incubated at 25 °C for 7 days in the dark. Thereafter, 220 µl (10^−5^M) of GR24 (a commonly used synthetic germination stimulant, Yoneyama et al. 2013) were added to the discs, and the petri dishes were resealed and kept in the dark for another 24 h.

### Cultivation in polyethylene bags

The PEB system of Parker & Dixon (1983), with the slight modification of Eizenberg et al. (2003) to tailor it to sunflower cultivation, was used for observing the sunflower root–*O. cumana* interaction, as follows. Sunflower seedlings at the cotyledon stage were placed on 25×10 cm glass microfiber filter papers (Whatman GF/A), which were then inserted into clear PEBs (35×10 cm), and allowed to grow in a growth chamber under controlled conditions (25 °C; 18 h light; 150– 200 µE m^−1^ s^−1^) for 10 days. Preconditioned *O. cumana* seeds were then carefully placed alongside the sunflower roots on the GF/A filter papers. Sterilized half-strength Hoagland nutrient solution (Hoagland and Arnon 1950) (5 ml) was supplied every day. Observations were carried out every 2–3 days with an electronic binocular microscope (Leica M80) to monitor seed germination, attachment and establishment or necrosis of the *O. cumana* seedlings.

### Histological analysis

Plant material for histological analysis was taken from the PEB system. ‘EMEK3’ and ‘D.Y3’ roots, along with the attached parts of *O. cumana* seedlings, were sampled 5 days post infection. In parallel, non-infected sunflower roots (control) were sampled. The sampled roots (with and without *O. cumana*) were fixed in FAA [5% formalin: 5% acetic acid: 90% alcohol (70%), v/v] for 24 h. Fixed samples were then dehydrated in an ethanol series (50, 70, 90, 95, 100%; 1–2 h each). After dehydration, the samples were infiltrated with a series of Histo-Clear:ethanol (1:3, 1:1, 3:1 ratio; 1 h each), cleared with Histo-Clear (xylene substitute) and embedded in paraffin. The samples were then cut into 13-µm sections with a rotary microtome (Leica RM2245, Leica Biosystems, Nussloch, Germany) and stained with safranin/fast green (Ruzin 1999).

### Grafting experiments

To assess the involvement of the shoot in the resistance mechanism, grafting experiments were conducted as follows: ‘EMEK3’ and ‘D.Y3’ seeds were sown in 2-l pots, and 14 days post emergence the stems of plants with two true leaves were cut above the cotyledons at a 45° angle. ‘EMEK3’ shoots were grafted onto ‘D.Y3’ rootstock and vice versa. The grafted sunflowers were kept in a closed chamber with 100% humidity at 25°C for 3 days. The plants were then transferred to a humid chamber (in which water was sprayed every 3 h for 10 s) for 7 days. Thereafter, the plants were gradually exposed to the atmosphere (10% for 2 days, 40% for 2 days, 70% for one day, 90% for one day and subsequently 100%), and when acclimated, they were planted in 2 liter pots inoculated with 15 ppm of *O. cumana* seeds. Self-grafted and non-grafted plants served as control.

### Statistical analysis

The experiments were carried out in 5 replications in a fully randomized design. The analysis of variance (ANOVA) was performed, and means were compared using Student’s t test (P < 0.05) in JMP PRO 12 software (v5.1; SAS Institute Inc., Cary, NC).

### Bulk construction and RNA extraction

Roots of PEB-cultured sunflowers of five resistant lines, five susceptible lines and the resistant cultivar, were collected on the day of infestation and at 5 days post infestation with *O. cumana* for both infected and control plants. Whole roots were ground in liquid nitrogen, and equal amounts of root tissue from each variety of the resistant and the susceptible lines were taken as R and S bulks. Total RNA was isolated from 27 samples (‘EMEK3’, R bulk and S bulk × 3 treatments/ sampling time × 3 replicates) using Spectrum™ Plant Total RNA Kit (Sigma-Aldrich) according to the manufacturer’s protocol. RNA quality and integrity were evaluated by Agilent TapeStation 2200 (Agilent Technologies).

### RNA sequencing and mapping

Libraries were prepared using the Genomics in-house protocol for mRNA-seq. Briefly, the polyA fraction (mRNA) was purified from 500 ng of total RNA, followed by fragmentation and the generation of double-stranded cDNA. Then, end repair, a base addition, adapter ligation and PCR amplification steps were performed. Libraries were evaluated by Qubit (Thermo Fisher Scientific) and TapeStation (Agilent). Sequencing libraries were constructed with barcodes to allow multiplexing of 27 samples in 2 lanes. Approximately 16-20 million single-end 60-bp reads were sequenced per sample on an Illumina HiSeq 2500 V4 instrument. The quality of the raw reads was evaluated using FastQC v.0.11.03 (Andrews 2010), followed by trimming and removal of low-quality reads using Trimmomatic v.036 (Bolger et al. 2014). Cleaned reads from each of the 27 libraries were then aligned to the *H. annuus* XRQ v1.0 reference genome (Badouin et al. 2017) using STAR v.2.5.2b (Alexander et al. 2013), and the level of expression of each gene in each library was estimated using RSEM v.1.2.31 (Li and Dewey 2011). Expression levels were normalized using the number of reads per kilobase per million reads mapped (RPKM) for each transcript.

### RNA-Seq data analysis

The RSEM output files were analyzed using R package DESeq2 (Love et al. 2014) for differential expression analysis. A pairwise comparisons test was performed between conditions in ‘EMEK3’, R bulk and S bulk. DEGs were considered as significant at FDR < 0.05 (Benjamini and Hochberg 1995). GO terms were obtained from the heliagene database for XRQ (https://www.heliagene.org/HanXRQ-SUNRISE/), and GO terms enrichment analysis was performed for significant DEGs compared to all other GO terms using the Blast2GO (v5.2.5) analysis tools (Götz et al 2008). Significantly over-represented GO terms were identified using Fisher’s exact test at significance level of FDR < 0.05. GO slim (Blast2GO tool) was performed to reduce the complexity of GO terms for functional analysis of annotated *H. annuus* genes.

## Abbreviations

C4H: Cinnamic acid 4-hydroxylase
DCL: Dehydrocostos Lactone
DEG: Differentially Expressed Genes
EREBP: Ethylene Responsive Element Binding Protein
ERF: Ethylene Responsive Factor
ETI: Effector Triggered Immunity
FPKM: Fragments Per Kilobase of exon per million fragments Mapped
HR: Hipper Sensitivity
LG: Linkage Group
LLR: Leucine Rich Repeat
MAMP: Microbe Associated Molecular Patterns
mRNA: messenger Ribonucleic acid
NBS: Nuclear Binding Site
PAL: Phenylalanine-ammonia Lyase
PAMP: Pathogen Associated Molecular Patterns
PEB: Polyethylene bag
POX: Peroxidase
PR: Pathogen Related
PTI: Pathogen Triggered Immunity
QTL: Quantitative Trait Locus
RAPD: Random Amplification of Polymorphic DNA
RNA: Ribonucleic acid
SNP: Single Nucleotide Polymorphism

## Acknowledgment

This project was funded by the Office of the Chief Scientist, Israel Ministry of Agriculture grant No. 132-1704-14.

## Author Contribution Statement

YD and HE conceived the idea and supervised the project overall. DS, YT, DP and HE designed the experiments. DS conducted experiments, analyzed data and wrote the manuscript. SH performed the differential expression analysis. All authors read and approved the manuscript.

---

**Supplementary Fig. 1.**
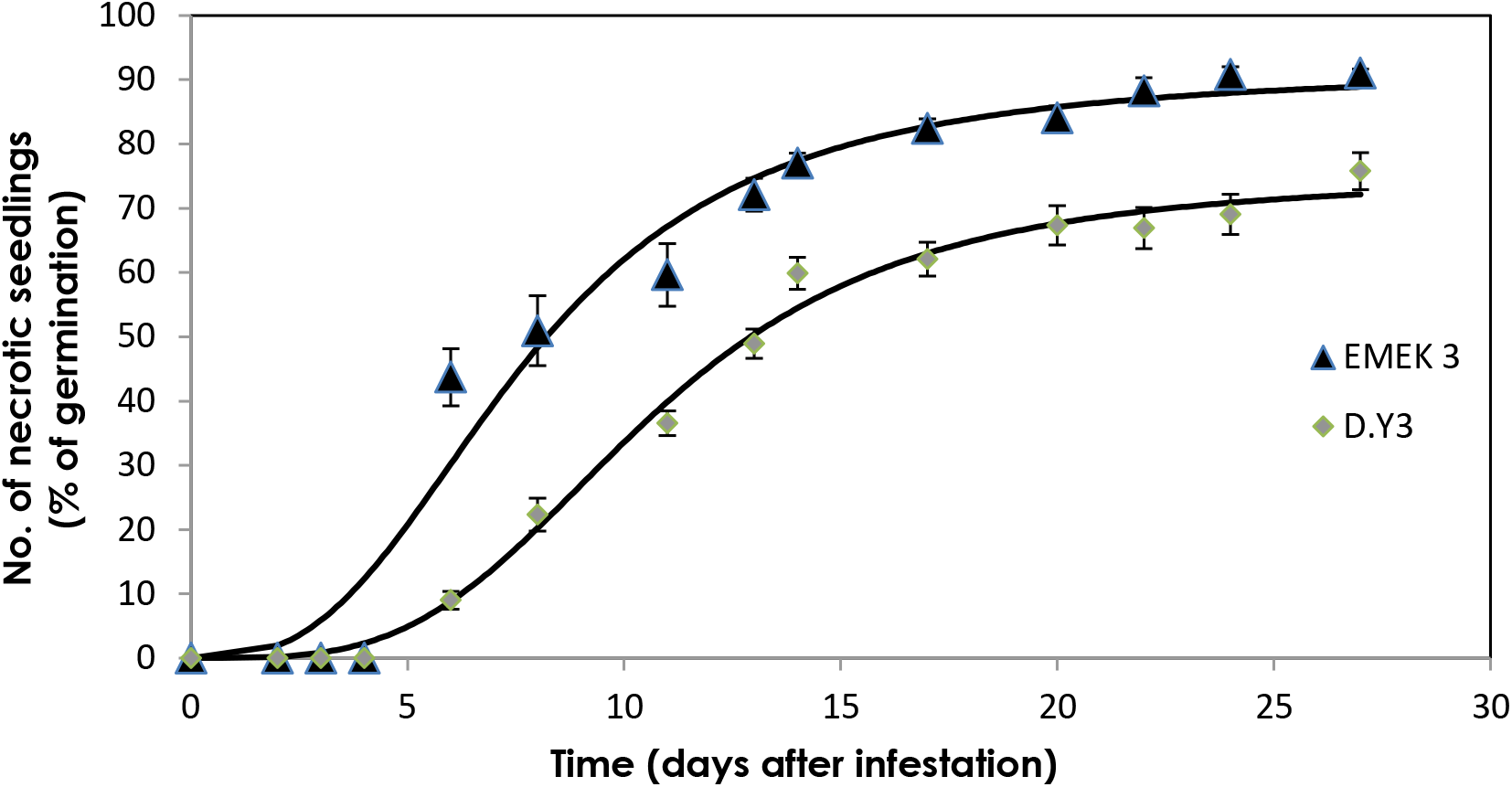
parasitism necrotic seedlings (% of germination) of *O. cumana* at the presence of resistant (EMEK3) and susceptible sunflower (D.Y3) varieties.

**Supplementary Fig. 2.**
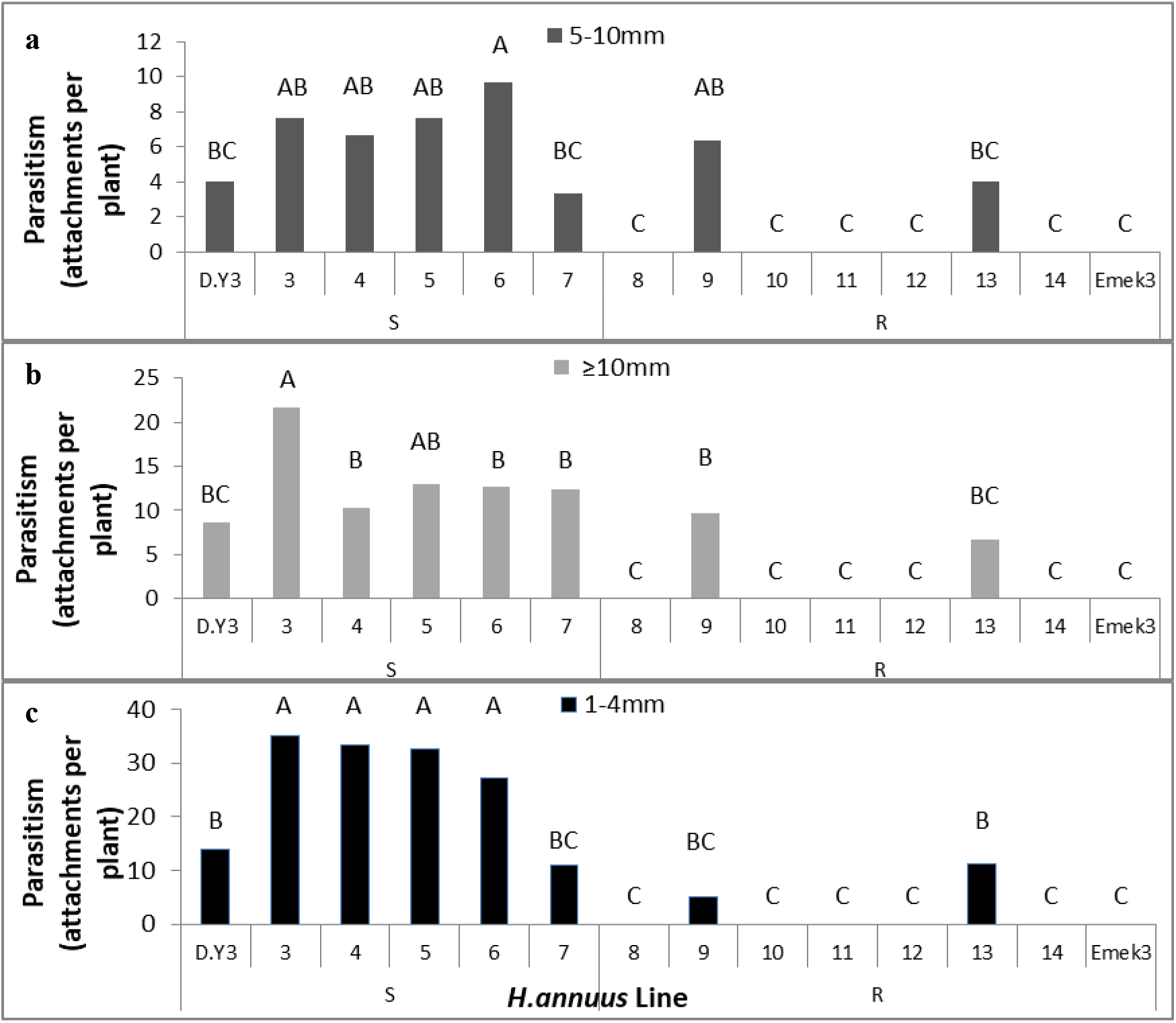
Number of parasitized *O. cumana* on the roots of the sunflower varieties ‘EMEK3’, ‘D.Y3’ and of the lines 3-14. (**a**) Number of 1-4mm tubercles; (**b**) Number of 5-10mm tubercles; (**c**) Number of tubercles larger than 10mm.

**Supplementary Table 1.** Number of reads of the 27 libraries.

| Library<br>ID | Index | # PF Reads |
| --- | --- | --- |
| ea1 | CATAGCGA | 9,131,020 |
| ea2 | CAGGAGCC | 8,964,291 |
| ea3 | TGTCGGAT | 9,513,800 |
| eb1 | ATTATGTT | 8,994,704 |
| eb2 | CCTACCAT | 9,225,671 |
| eb3 | TACTTAGC | 9,366,030 |
| ec1 | GAAGAAGT | 8,940,419 |
| ec2 | AGGATCTA | 8,852,028 |
| ec3 | GACAGTAA | 8,973,442 |
| ra1 | GGTCCAGA | 8,500,951 |
| ra2 | GTATAACA | 8,507,016 |
| ra3 | TTCGCTGA | 8,074,534 |
| rb1 | AACTTGAC | 8,145,121 |
| rb2 | CACATCCT | 8,697,047 |
| rb3 | TCGGAATG | 8,593,227 |
| rc1 | AAGGATGT | 8,098,075 |
| rc2 | GCACACGA | 9,199,815 |
| rc3 | TCTGGCGA | 9,807,335 |
| sa1 | CTACCAGG | 7,868,629 |
| sa2 | CATGCTTA | 7,430,950 |
| sa3 | GCACATCT | 8,043,633 |
| sb1 | TGCTCGAC | 8,162,395 |
| sb2 | AGCAATTC | 9,304,152 |
| sb3 | AGTTGCTT | 8,829,022 |
| sc1 | CCAGTTAG | 8,769,869 |
| sc2 | TTGAGCCT | 7,639,542 |
| sc3 | ACCAACTG | 7,886,680 |

